# Understanding the bacterial imbalance in Hidradenitis Suppurativa patients: Insights into microbial community shifts and colonization by opportunistic pathogens

**DOI:** 10.1101/2023.11.12.566737

**Authors:** Lene Bens, Tine Vanhoutvin, Alison Kerremans, Daan Jansen, Melissa Depypere, Tom Hillary, Séverine Vermeire, Sabrina I. Green, Jelle Matthijnssens, João Sabino, Rob Lavigne, An Van Laethem, Jeroen Wagemans

## Abstract

Patients suffering from hidradenitis suppurativa (HS) develop painful skin lesions, significantly decreasing their quality of life. This chronic disease is triggered by plugged hair follicles resulting in an aberrant immune response, skin microbiome imbalance and secondary bacterial colonization. As a result, a diversity of treatment options are currently applied, including antibiotics, biologicals like adalimumab and surgery, which often provide only short-term relief. Alternative strategies, like phage therapy, have been proposed but identification of the target bacterium is key. Therefore, a spatial and longitudinal analysis was performed on skin swabs of lesions from 39 HS patients and 18 healthy controls, leading to a total collection of 108 lesional samples and 35 control samples at different time points and locations throughout the body. Samples were subjected to 16S rRNA community analysis, as well as bacterial isolation using aerobic and anaerobic culturing in combination with MALDI-TOF. Our data demonstrate that the bacterial community present in lesions of patients with HS is out of balance compared to healthy individuals, in which the niche of *Staphylococcus* and *Corynebacterium* is taken over by *Escherichia*-*Shigella*. Overall, three bacterial community profiles of HS lesions and one of healthy individuals could be distinguished. Although the overall bacterial composition was not associated with the disease severity defined by the Hurley classification system, lesions often become colonized with opportunistic pathogens including *Staphylococcus aureus* and *Pseudomonas aeruginosa* at increasing disease severities. Furthermore, patients with a concurrent IBD diagnosis did not reveal a significantly different bacterial skin community.

## Introduction

Hidradenitis suppurativa (HS) or *acne inversa* is a chronic inflammatory skin disease characterized by plugging of the hair follicles which lead to the formation of painful noduli, abscesses and fistulas. This skin disease with unknown cause affects 0.03% to 4% of the global population^1–3^. Moreover, it is associated with comorbidities like inflammatory bowel disease (IBD)^4,5^. Development can be triggered by a variety of factors, including hormonal, genetic, physical and environmental factors. However, the immune system as well as the skin microbiome are thought to play an important role in disease emergence^1,6^. Previous research showed that the microbiome of HS patients differs in both lesional and non-lesional skin when compared to healthy individuals^7^. This microbial imbalance potentially leads to an aberrant immune response, finally resulting in clogging of the follicles. Interestingly, it is also hypothesised that a misdirected immune response causes this bacterial imbalance. Nevertheless, an interplay of the immune response and microbiome presumably lies at the basis of this disease. Skin lesions manifest primarily at the axillary, inguinal and anogenital regions of the body and can become colonized by opportunistic bacteria which again boost inflammation, starting a vicious cycle ^2,8^.

Antibiotics, biologicals like adalimumab and surgery are a number of treatment options that are currently applied for HS^9^. However, these treatments often only provide short-term relief of disease symptoms in case the patient responded to the treatment in the first place^10,11^. As a complementary strategy, we recently proposed phage therapy, but its implementation depends on the identification of the target bacterium^12^. Previously, both culturing and 16S rRNA profiling have been carried out in which the presence of both commensal bacterial species as well as primary pathogens were observed. Although *Staphylococcus*, *Streptococcus* and *Corynebacterium* were regularly identified in HS lesions, no specific bacterial pattern could be associated with HS^12^.

Here, we examine the bacterial composition from lesions of 39 HS patients with varying disease severity and 18 healthy individuals at multiple time points and locations across the body. Within this cohort, patients suffering from both HS and IBD have also been included to examine whether distinct microbiomes can be observed between HS patients with and without concurrent IBD diagnosis. Moreover, a parallel study focuses on the virome present in these HS lesions, revealing a higher abundance of *Corynebacterium*- and *Staphylococcus*-infecting phages in healthy individuals^13^.

## Material and methods

### Study design

This study was carried out under the approval of the UZ Leuven (Belgium) ethical committee (s62301). All individuals signed an informed consent to participate and sampling was carried out under the DALIHS biobank.

### Patient and healthy control selection

Patients attending the UZ Leuven clinic (Leuven, Belgium) with a history of HS were included and disease severity was assessed using the Hurley classification^14,15^. Patients were selected to ensure at least ten patients for each of the three Hurley stages. Furthermore, patients with both HS and IBD were included. As controls, healthy individuals without dermatological disorders were chosen to match the gender ratio of the patient cohort. For each individual, the following data were included: gender, age, Hurley stage (Hurley I, Hurley II, Hurley III), presence of IBD (yes, no), smoking status (yes, no, past), ethnicity, BMI category (underweight: BMI < 18.5 kg/m^2^, normal: 18.5 kg/m^2^ ≤ BMI ≤ 25 kg/m^2^, overweight: 25 kg/m^2^ < BMI ≤ 30 kg/m^2^, obese: 30 kg/m^2^ < BMI) and living place (Supplementary Table S1).

### Sample collection

At multiple time points over one year, three replicate skin swabs were taken from one to three regions of the patient where HS lesions were present. For healthy individuals, the axillar and groin regions were sampled. For each HS sample, the activity of the lesion (nodulus, abscess, fistula, remission after deroofing, remission), the sampling location (axilla, abdomen, buttocks, chest, groin & genitals) and treatment at time of sampling (antibiotic, biological, intralesional corticosteroid, hormone suppressant, metformin, methotrexate, only local treatment, vitamin A acid, zinc supplement) was noted. All patients following a treatment also included a local treatment of antiseptic wash, and antiseptic and keratolytic ointments. The liquid of one swab was stored at −80°C for 16S rRNA profiling.

### Bacterial culturing

For aerobic growth, skin swabs were streaked onto blood culture plates and incubated for one week at 37°C. For anaerobic growth, a tube with thioglycolate broth was inoculated with the skin swab and incubated for one week at 37°C. Next, the ‘thio’ broth was streaked onto anaerobic blood plates and incubated anaerobically in an anaerobic growth chamber (90% N_2_, 5% CO_2_, 5% H_2_)for a week at 37°C. Pure isolates were purified by re-culturing single colonies on fresh blood plates.

### Taxonomic classification of bacterial isolates

Bacterial isolates were taxonomically classified by using matrix-assisted laser desorption/ionization time-of-flight (MALDI-TOF) mass spectrometry (Bruker UK Ltd, MBT compass library Revision L). For a limited number of isolates, the species or even genus/family could not be determined with the available MALDI-TOF database. In those cases, the 16S rRNA gene was amplified with PCR (universal primers 27F - AGAGTTTGATCMTGGCTCAG and 1492R - CGGTTACCTTGTTACGACTT) and sequenced with Sanger sequencing to determine the genus by blasting the sequenced 16S rRNA gene against the 16S rRNA database of NCBI (consulted on 03.10.2022)^16^.

### Statistical analysis of culturing data

Statistical analysis was performed with JMP Pro 16.0.0 at significance level 0.05. Associations between the number of isolated strains per sample and the meta-data was assessed with a Kruskal-Wallis test. A Pearson *X*^2^ test was used to examine the association between nominal data of samples: Hurley stage, region of sampling and presence/absence of bacterial genus. When the sample count was below five for 20% of the nominal data combinations, a Fisher’s exact test was performed.

### Bacterial composition analyses using 16S rRNA gene amplification & sequencing

The DNA of the swabs was extracted with the DNeasy UltraClean Microbial kit (Qiagen) according to the manufacturers protocol, although the elution step was performed with 20 µL buffer instead of 50 µL. In total, 108 samples of HS patients and 35 samples of healthy controls were processed. After DNA isolation, the V4 region of the 16S rRNA gene was amplified with the primers described by Pichler *et al.* which enabled dual-index barcoding for the Illumina MiniSeq platform^17^. Next, DNA was purified with AMPure beads and the DNA concentrations were measured with the Qubit fluorometer. After pooling the samples together in equimolar concentrations in resuspension buffer (Illumina), the MiniSeq denaturation protocol A (Illumina) was followed. Samples were sequenced with the Illumina MiniSeq device using the Illumina MiniSeq Mid output 300 cycles kit (2×150 bp, paired-end, Illumina). Demultiplexing and trimming of the reads was performed using the Local Run Manager (Illumina).

### Taxonomic classification of 16S rRNA sequencing

Processing of the sequencing reads into operational taxonomic units (OTUs) and taxonomic classification was achieved using QIIME2 version 2022.8.0^18^. First, the quality profile of 10,000 random sequences was determined with *demux summarize* which was used to determine the parameters of the read processing^18^. Denoising and clustering of the data into OTUs was performed using dada2 with truncation lengths set at 150 bp for both forward and reverse reads^18–20^. A SILVA database (version 138.1) was created by the machine learning-based classification method and used in QIIME2 with *feature-classifier classify-sklearn* to assign taxonomy to the features^18–27^. Then normalization of the data was carried out by scaling with ranked subsampling (SRS), for which the C_min_ (12,225) was determined by the Shiny app^28^, leading to the loss of five HS and eight control samples.

### Diversity analysis

16S rRNA analysis, statistical tests and visualization were performed at the genus-level with R version 4.2.2^29^ by making use of the *qiime2R*^30^, *phyloseq*^31^ and *ggplot2*^32^ packages. When samples were pooled, the mean abundance was used. Differentially abundant genera were identified with *ALDEx2*^33^ (Wilcoxon test, Benjamini-Hochberg (BH) corrected). Alpha-diversity, including observed taxa and Shannon diversity, was determined with *phyloseq*. Median differences between two groups were tested with a Mann-Whitney U test. When the difference between three or more groups was examined, a Kruskal-Wallis test was performed. Beta-diversity (Bray-Curtis) was also calculated with *phyloseq*. Differences in β-diversity were assessed with the nonparametric Adonis test (PERMANOVA) in *vegan*. Bray-Curtis dissimilarity ordination was visualized with a principal-coordinate analysis (PCoA).

### Metadata contribution to microbial composition

The contribution of metadata variables on the microbial composition was determined by distance-based redundancy analysis (dbRDA). The independent contribution was determined with the capscale function of *vegan*^34^, for which a BH correction for multiple testing was applied. Variables that showed a significant contribution to the ordination were further examined in a stepwise dbRDA (ordiR2step of *vegan*) to determine the cumulative contribution. The effect of the metadata on and the taxonomic association strength with the first two axes of the ordination was determined with the envfit function of *vegan* after which they were plotted on the PCoA plot. For all statistical tests, a significance level of 0.05 was used.

### Hierarchical clustering

A hierarchical clustering heatmap was constructed at the genus-level with R version 4.2.2^29^ with *pheatmap*^35^ with log-scale visualisation of abundance. Association of metadata to clustering of samples was examined with JMP Pro 16.0.0 at significance level 0.05, with a Pearson *X*^2^ test. When 20% of examined combinations had an expected sample count below five, a Fisher’s exact test was carried out. In case a variable with more two levels had a significant association, a correspondence analysis biplot was used to identify strong associations.

### Data availability

Raw sequencing data of 16S rRNA amplicons is available from NCBI under BioProject PRJNA1032850.

## Results

### Patient cohort and sampling

The HS patient cohort, as shown in Table 1, consisted of 39 individuals, aged 19 to 64 years (mean of 38 years ± 12.2), including 13 (33.3%) males and 26 (66.7%) females which is in accordance with the male to female ratio of HS (1:3.3) described in Western countries^36^. The majority (61.5%) is currently smoking with an additional 15.4% individuals who smoked in the past. Patients with different disease severities were selected and classified according to Hurley stages, i.e., 10 patients in Hurley stage I (25.6%), 19 in Hurley stage II (48.7%) and 10 in Hurley stage III (25.6%). As this disease is linked to comorbidities like IBD, also patients suffering from both HS and IBD (n = 12, 30.8%) were included to compare the bacterial community of HS patients diagnosed with and without IBD. Additional meta-data of patients and samples are shown in Supplementary Table S1 and S2, respectively.

**Table 1:**
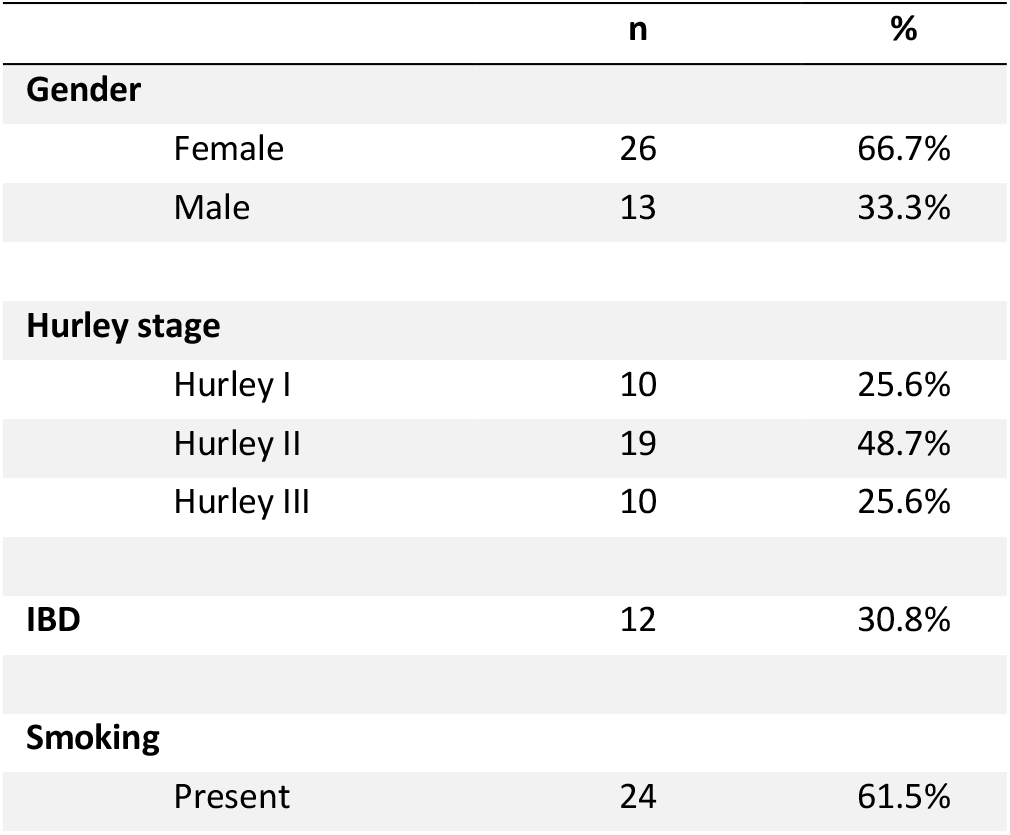
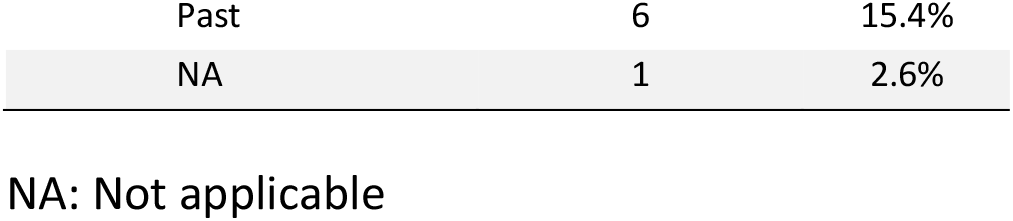
HS patient cohort characteristics.

|  | n | % |
| --- | --- | --- |
| **Gender** |  |  |
| Female | 26 | 66.7% |
| Male | 13 | 33.3% |
| **Hurley stage** |  |  |
| Hurley I | 10 | 25.6% |
| Hurley II | 19 | 48.7% |
| Hurley III | 10 | 25.6% |
| **IBD** | 12 | 30.8% |
| **Smoking** |  |  |
| Present | 24 | 61.5% |
| Past | 6 | 15.4% |
| NA | 1 | 2.6% |

NA: Not applicable

Swabs of skin lesions were taken longitudinally and spatially, leading to a total collection of 108 samples at different time points and at a diversity of locations across the human body, with the majority isolated from the typical zones where HS occurs: the axilla (n = 37), groin and genitals (n = 47), and buttocks (n = 13). Only a minority of samples were located at the chest (n = 8) and abdomen (n = 3). Three regions were defined: 1) the axilla, 2) chest & abdomen, and 3) groin, genitals & buttocks, and no significant association between Hurley stage and region of sampling was detected (Fisher’s exact test, p = 0.1746).

### Bacterial culturing does not reveal a single disease-causing pathogen

In total, 103 samples were processed for aerobic culturing, whereas 45 available samples were processed anaerobically. An overview of the isolated species and frequency of isolation is provided in Table 2, whereas the isolated species per sample is shown in Supplementary table S3. Polymicrobial growth on plate was visible for the majority of samples, whereas only two swabs showed no bacterial growth both under aerobic and anaerobic conditions. Overall, 296 different isolates were cultured from which only eight obligate anaerobes. Four of these, *Peptoniphilus*, *Cutibacterium*, *Peptostreptococcus* and *Finegoldia magna*, were also detected in previous HS studies^37–41^. However, since only one strain was detected of each genus across all samples, it is unlikely that these bacteria are associated with HS in our patient cohort. Bacteria belonging to the *Staphylococcus* genus were isolated most frequently from HS lesions. Coagulase-negative staphylococci (*S. epidermidis, S. haemolyticus* and *S. lugdunensis*), as well as coagulase-positive staphylococci (*S. aureus*) were identified. The second most prevalent genus present was *Corynebacterium*, also part of the known healthy human skin microbiota^42,43^. In addition, *Enterococcus*, *Streptococcus*, *Escherichia* and *Proteus* were detected regularly (n>10), which is in accordance with other culture-based studies^12^.

**Table 2:**
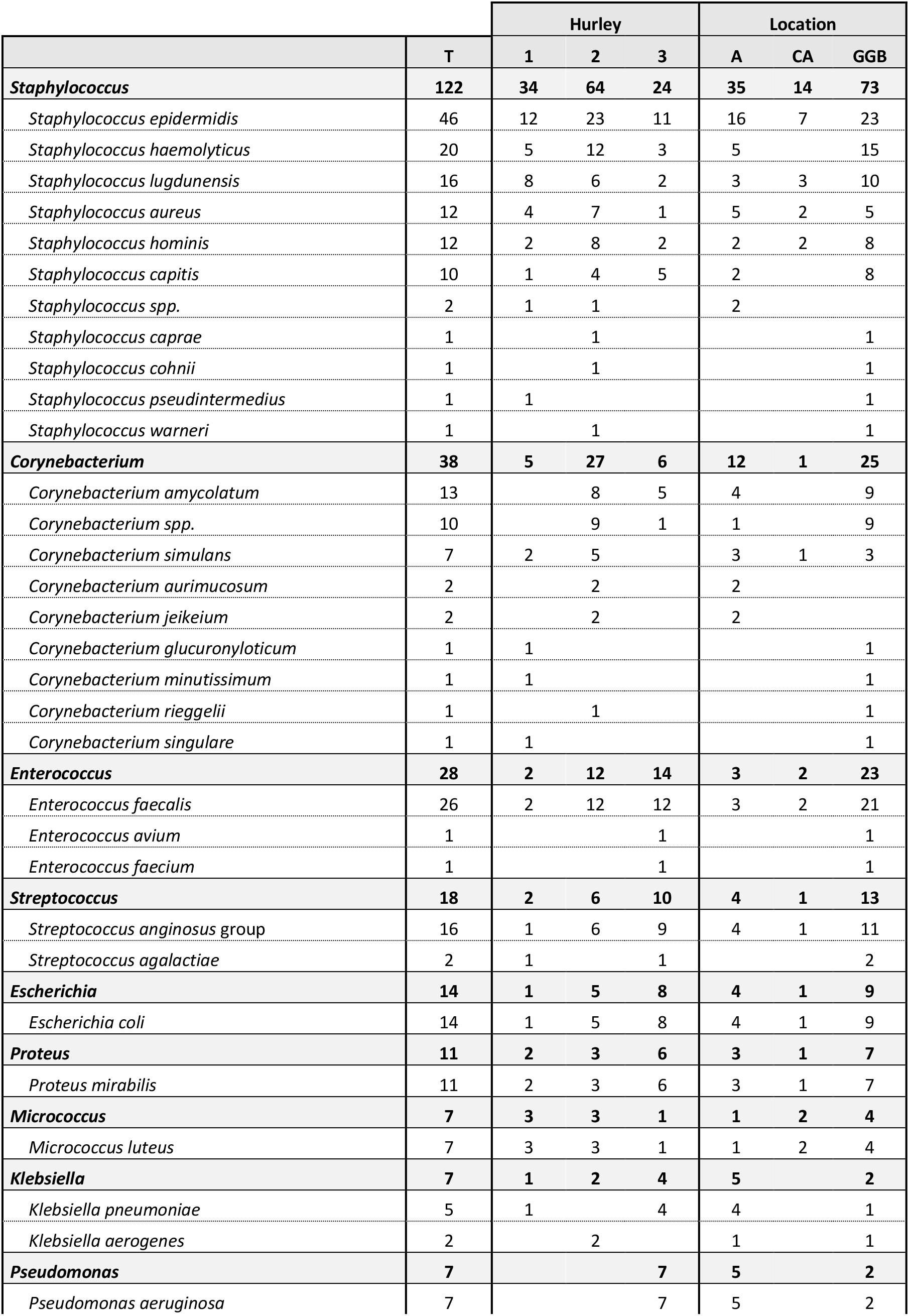
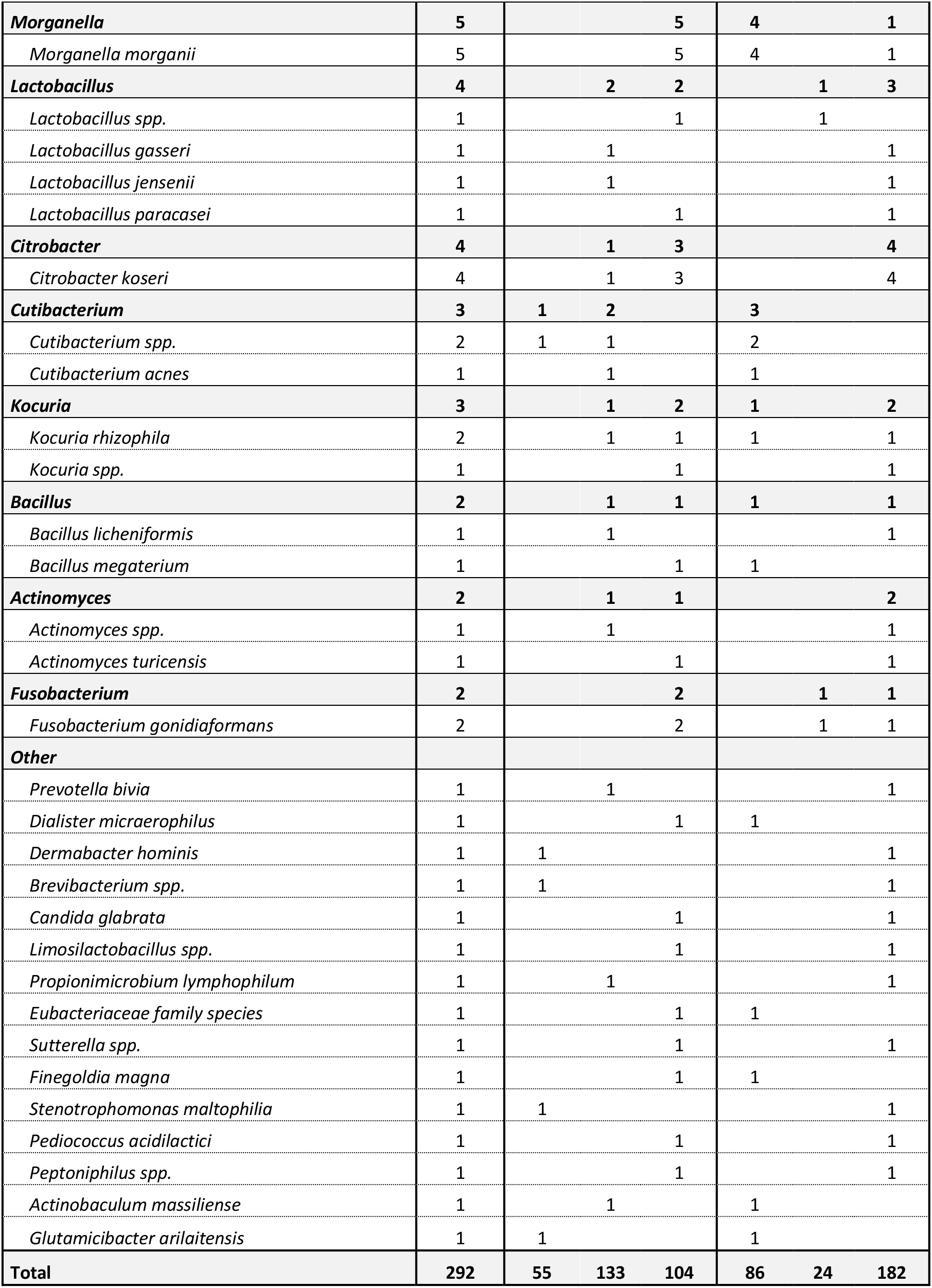
Number of isolates cultured per species and genus. Both the total number of isolates (T), as well as the number for each hurley stage and location (A: Axillae, CA: chest & abdomen, GGB: groin, genitals & buttocks) of sampling is shown. Note: per sample, only one isolate for each identified species was counted.

When samples are subdivided based on the Hurley stage of the patient, no association between Hurley stage and the number of bacterial species isolated could be observed (Kruskal-Wallis, p = 0.2380). However, there was a significant association with *Staphylococcus* (Pearson *X*^2^, p = 0.0048) being isolated less frequently with increasing Hurley stage, whereas *Enterococcus* (Pearson *X*^2^, p = 0.0235) and *Streptococcus* (Pearson *X*^2^, p = 0.0343) were isolated more frequently with increasing Hurley stage. At the species level, this translates in frequency differences for *Staphylococcus lugdunensis* (Fisher exact test, p = 0.0068), *Enterococcus faecalis* (Pearson *X*^2^, p = 0.0496) and *Streptococcus anginosus* (Fisher exact test, p = 0.0377). Furthermore, *Pseudomonas aeruginosa* and *Morganella morganii* were only isolated in samples of Hurley stage 3, in seven and five samples respectively.

When stratifying the bacterial species according to the localization of the lesions, no association between the number of bacterial isolates and region was observed (Kruskal-Wallis, p = 0.5100). Nevertheless, isolation of the genus *Enterococcus* could be linked to the localization. Indeed, this genus was isolated more frequently at the groin, genitals and buttocks (Pearson *X*^2^, p = 0.0120). As no association between Hurley stage and region of sampling exists, the more frequent isolation of *Enterococcus* at this location is not due to a dependency with the disease severity. Nevertheless, this higher frequency of isolating *Enterococcus* at this region compared to the others is not a surprise as species of this genus are part of the vaginal microbiome and most patients were female. In addition, *Enterococcus* is a subdominant genus present in the gastro-intestinal tract^44^.

### 16S rRNA profiling identifies a distinct bacterial skin community in HS lesions

16S rRNA profiling was carried out on a total of 144 samples, including 108 lesional samples of HS patients and 35 samples of healthy controls. After normalization, 103 HS samples and 27 control had a sufficient read count to be used for further analyses. For these, the taxonomic distribution at the genus level is shown in Supplementary Figure S1. Both the HS and control group showed a similar richness (Observed α-diversity, Mann-Whitney U, p = 0.2214), but the Shannon diversity of the bacteria is different (Shannon α-diversity, Mann-Whitney U, p = 0.0001) as the bacterial distribution in HS samples is more unequal (Figure 1A). Differential abundance analysis (Figure 1B) revealed ten bacterial genera to be significantly differentially abundant (ALDEx2, Benjamini-Hochberg corrected p < 0.05). *Escherichia*-*Shigella*, *Streptococcus*, *Pseudomonas*, *Porphyromonas* and *Bradyrhizobium* species are more abundant in lesions from HS patients compared to the healthy skin microbiome, whereas *Corynebacterium, Staphylococcus, Dermabacter, Paracoccus* and *Gluconobacter* are less abundant. The genera *Escherichia-Shigella* were observed most in HS samples with a mean relative abundance (mRA) of 0.38, whereas in healthy controls *Corynebacterium* and *Staphylococcus* were observed the most, with a mRA of 0.43 and 0.33, respectively. Note that the other differentially abundant genera have a mRA below 0.1.

**Figure 1.**
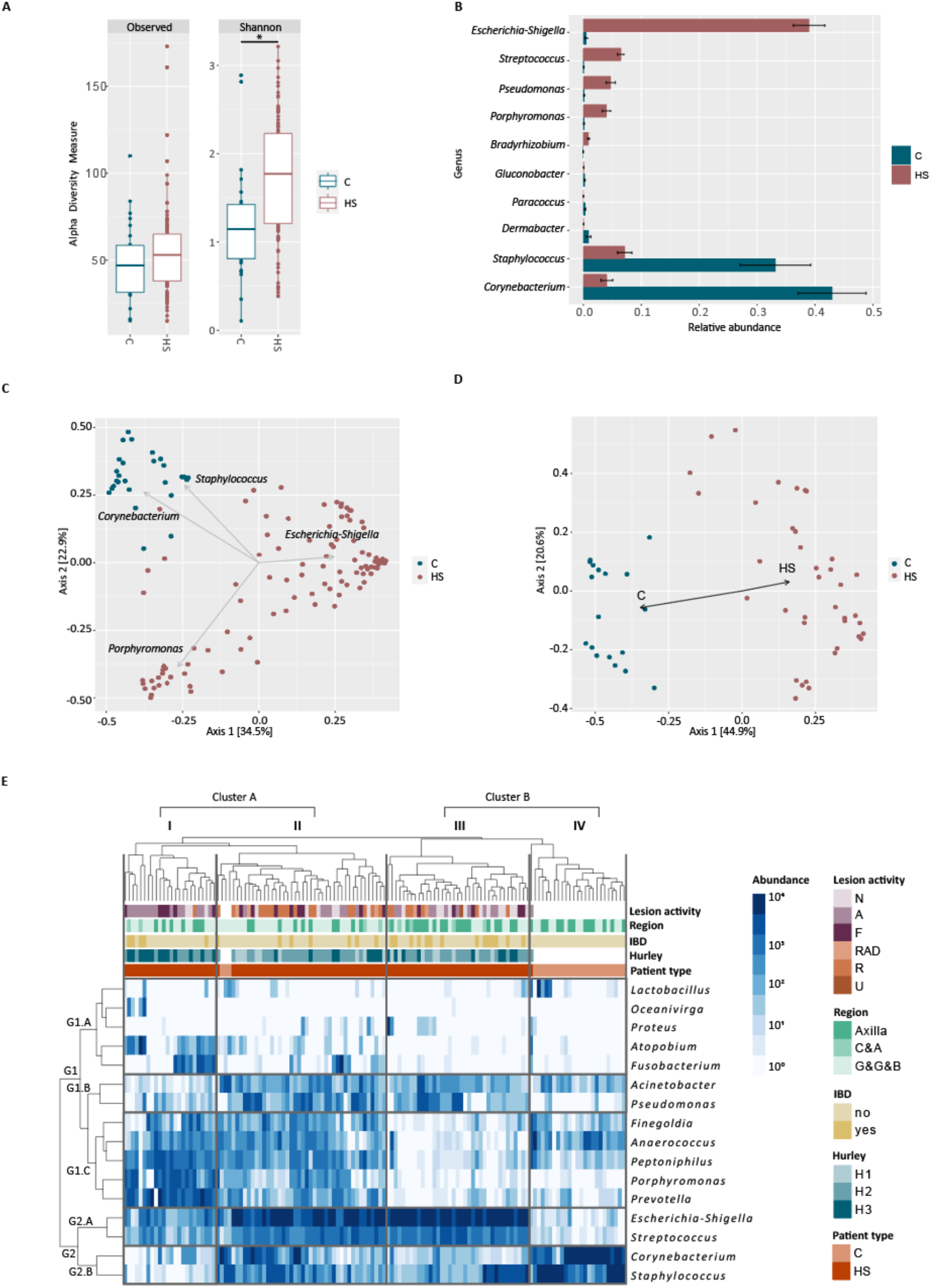
Control (C) and HS. **A) α-diversity.** Both the observed number of genera and Shannon α-diversity is shown (*p-value = 0.00018). **B) Differential abundance analysis (DAA)**. The mean relative abundance of genera identified as significantly differentially abundant are shown. **C) PCoA of Bray-Curtis dissimilarity (β-diversity).** The differences in microbiota composition is visualized as a distance between the points. The strength of the taxa association to the ordination is indicated with arrows. **D) PCoA of Bray-Curtis dissimilarity (β-diversity), samples pooled per individual.** The differences in microbiota composition are visualized as a distance between the points. The strength of the disease status to the ordination is indicated with arrows. **E) Hierarchical clustering.** The relative abundance of the bacterial genus is shown in logarithmic scale. Only genera with a minimal abundance of 15% in one of the samples were included. For each sample, the lesion activity and region of sampling, and smoking status, BMI category, IBD presence, hurley stage and presence of HS of the patient is indicated.

The community-level differences, or β-diversity, was examined with the Bray-Curtis dissimilarity and ordination was visualized with a two-dimensional principal-coordinate analysis (PCoA) (Figure 1C). On this PCoA, 56.5% of microbiome variation is visualised and separate grouping of the control samples and HS samples can visually be observed. Indeed, the Bray-Curtis β-diversity of the control and HS group is significantly different (PERMANOVA, p-value = 0.001), meaning the HS and control samples have distinct bacterial communities. From the differentially abundant genera (Figure 1B), *Corynebacterium*, *Staphylococcus*, *Escherichia*-*Shigella* and *Porphyromonas* showed a significant association with the ordination, with the first two being associated with control samples and the latter two with HS. This visually shows that two groups in the HS population can be distinguished based on a higher abundance of either *Escherichia-Shigella* or *Porphyromonas*.

Univariate distance-based redundancy analysis (dbRDA, Bray-Curtis) on the samples pooled per individual showed that disease status (R^2^adj = 0.380, BH corrected p = 0.003), living place category (R^2^adj = 0.22, BH corrected p = 0.003), smoking (R^2^adj = 0.075, BH corrected p = 0.02) and age (R^2^adj = 0.041, BH corrected p = 0.021) have a significant effect on the bacterial composition. However, only disease status, thus HS or control, has a significant effect in multivariate dbRDA, showing that only the presence or absence of HS has an independent effect on microbial composition, explaining 38% of variation (Figure 1D). To assess whether the location of sampling is associated with the composition of bacterial genera, the same analysis was carried out on the individual samples, but did not indicate any significant effect of sampling location on the microbial composition (data not shown). This suggests that HS patients have a lesional microbiome that is distinct from the healthy skin microbiome, independent of the location on the human body.

### Three HS lesional bacterial profiles

Hierarchical clustering was carried out to examine sample clustering based on the distribution of bacterial genera. Only genera with a minimal abundance of 15% in at least one of the samples were included for this analysis (Figure 1E). Clustering of different bacteria (horizontally) based on their similarities in their abundance is visualised on the left side. In here, two main clusters (G1, G2) and five subgroups are distinguished. Group G1.A comprises five genera, *Lactobacillus*, *Oceanivirga*, *Proteus*, *Atopobium* and *Fusobacterium*, with each a high abundance in only one or a few samples, indicating that there is probably no association of these genera with the microbiome of healthy or diseased samples. Furthermore, *Pseudomonas* and *Acinetobacter* group together in G1.B, whereas a third group within cluster G1 only comprises obligate anaerobic genera *Finegoldia*, *Anaerococcus*, *Peptoniphilus*, *Porphyromonas* and *Prevotella* (G1.C). For cluster G2, there is a subgrouping of *Escherichia-Shigella* and *Streptococcus* in G2.A, and *Corynebacterium* and *Staphylococcus* in G2.B.

Samples (columns) are clustered together according to their microbial composition, leading to the identification of two cluster (A and B), containing four bacterial community profiles (I-IV; Supplementary table S4). Samples in Cluster A are characterized by a difference in abundance of obligate anaerobic bacterial genera (G1.C). Inside cluster A – which mainly contains HS samples (95.6%) - two distinct microbiome profiles can be distinguished. Profile I has a medium relative abundance of *Escherichia*-*Shigella* and *Streptococcus* (G2.A), and a low relative abundance of *Staphylococcus* and *Corynebacterium* (G2.B), whereas, profile II has a high relative abundance of *Escherichia*-*Shigella* and *Streptococcus*, and a medium to low abundance of *Staphylococcus* and *Corynebacterium*. Cluster B on the other hand has a low abundance of obligate anaerobes and contains both HS and control samples. However, most of the HS samples can be subdivided in the third HS microbiome profile (III), characterized by a high abundance of *Escherichia*-*Shigella* and *Streptococcus*, which are low abundant in microbiome profile IV, which primarily consists of control samples (96%). This microbiome profile of healthy samples is characterized by a low abundance of obligate anaerobes, *Escherichia*-*Shigella* and *Streptococcus*, and a high abundance of *Corynebacterium* and *Staphylococcus*.

When assessing disease severity, an association between Hurley stage and clustering of the samples into microbiome profiles was observed (Pearson *X*^2^, p = 0.0380). Specifically in profile I, no Hurley stage I samples were present. Furthermore, the activity of the lesions, referring to the presence of noduli, abscesses or fistulas, also correlated with the profiles (Pearson *X*^2^, p = 0.0058). The activity of the lesions was independent of the Hurley stage (Pearson *X*^2^, p = 0.4334). In case the patient had only noduli, this was strongly correlated with profile III. As this type of lesion is less deep and no puss is drained, it explains the low abundance of obligate anaerobes due to more superficial sampling. Surprisingly, samples of lesions in remission, referring to locations with previously active lesions that completely healed at the time of sampling, were associated with profile II, instead of the HS profile III, which is closer to the healthy profile. However, profile II has a medium abundance of *Staphylococcus* and *Corynebacterium*, which are two genera of which the healthy microbiome is mainly constituted, potentially explaining why lesions in rest have this profile.

### Hurley staging is not correlated with distinct bacterial compositions

The Hurley stages are often used by physicians to grade the extent of HS severity. In view of bacterial-targeted therapy, it is important to know whether there is a different microbial composition in patients with a different disease severity. For this purpose, only the HS samples were selected and analysis of the microbial composition was carried out on this subset. With DAA, it was determined that no genera are differently abundant in patients with different disease severities. In addition, no difference in microbial richness and evenness is present as no significant difference in observed taxa and Shannon diversity is observed (Figure 2A).

**Figure 2.**
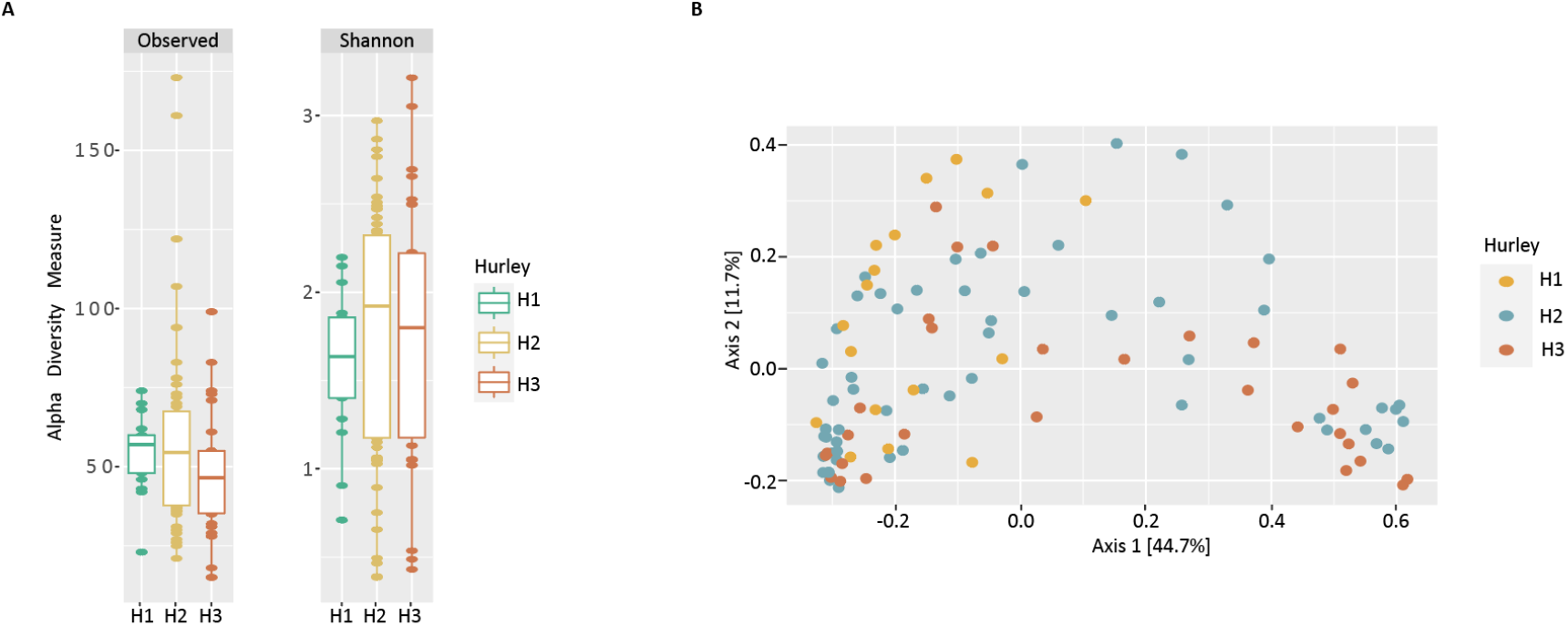
**A) α-diversity for different hurley stages**. Both the observed number of genera and Shannon α-diversity is shown for the Hurley 1 (H1), Hurley 2 (H2) and Hurley 3 (H3) group. **B) PCoA of Bray-Curtis dissimilarity of HS samples.** Samples are indicated as dots, including Hurley 1 (H1, yellow), Hurley 2 (H2, blue) and Hurley 3 (H3, red). The differences between microbiota composition is visualized as the distance between points.

The Bray-Curtis dissimilarity distances of the HS samples were calculated and 56.4 % of data variation is shown on the ordination in Figure 2B. Univariate distance-based redundancy analysis (dbRDA, Bray-Curtis) did not reveal any association of the microbial composition with patient-specific variables (gender, age, ethnicity, smoking, residence, Hurley stage, presence of IBD, obesity and lesion activity).

## Discussion

These data illustrate that the microbiome of HS lesions significantly differs from that of healthy controls, consistent with previous studies^7,45–47^. Although both groups have a similar bacterial richness, the distribution of the bacterial taxa is significantly different, as illustrated by the higher Shannon diversity index (fig. 1A). This demonstrates that HS lesions have a bacterial community that is out of balance, in which *Escherichia*-*Shigella* are the main key genera that take over the niche from *Staphylococcus* and *Corynebacterium* in our patient cohort (fig. 1B). However, these latter two genera were cultured most from HS swabs, whereas *E. coli* was only cultured in 14 samples, showing a discrepancy between the results of culturing and 16S rRNA profiling. This was expected and the principal reason why both culture-based and sequence-based identification techniques were performed as complementary approaches. Indeed, while culturing fails to capture entire spectrum of bacteria present, 16S rRNA profiling extends the identifications to viable but not culturable (VBNC) bacteria^48–50^. Nevertheless, the difference in abundance of *Staphylococcus* and *Corynebacterium* between these groups is also supported by the identification of phages predicted to target these bacteria, as shown by Jansen *et al*.^13^. They show that samples from healthy controls are associated with the presence of temperate phages infecting these two genera, confirming their presence and potentially increasing their fitness and stabilizing the skin microbiome. Taken together, this may explain their lower abundance in lesion samples. It is also important to highlight that 16S rRNA analysis only allows differentiation of bacterial taxa down to the genus level^51^. As a result, the identification of bacterial (sub)species can be difficult, although this might be important from a treatment point of view. For example, the genus *Staphylococcus* contains both commensal species such as *S. epidermidis* and pathogenic species such as *S. aureus*, a pathogen responsible for numerous skin infections^52^. Although the genus *Staphylococcus* is less abundant in HS lesions, it remains important to consider pathogenic *Staphylococcus* species as a bacterial target for alternative antimicrobial treatments, such as phage therapy^12^.

Both differential abundance analysis and ordination of the beta-diversity showed that *Escherichia-shigella* is the principal bacterium taking over the niche in HS patients. Although present in lower abundance, *Porphyromonas* was also identified with a higher abundance in HS samples. Surprisingly, the identification of *Escherichia-Shigella* as a potential key player in HS pathogenesis was not reported previously, whereas the higher abundance of *Porphyromonas* was reported in multiple studies^7,45–47,53^. Nevertheless, Hispan *et al.* found a significant higher detection of bacterial DNA in peripheral blood samples of HS patients, which were classified as Gram-negative bacillus DNA, especially *E. coli,* supporting the potential importance of this bacterial species in HS pathology^54^.

In total, four bacterial profiles, including three HS-dominated profiles and one healthy control dominated profile, were identified based on the presence or absence of a specific bacterial taxa (Figure 1E). Type I is characterized by a high abundance of obligate anaerobes, including *Porphyromonas*. A high abundance of *Escherichia-Shigella* and *Streptococcus* is present in type III, while a high abundance of both anaerobes, *Escherichia-Shigella* and *Streptococcus* is present in profile II. By contrast, healthy samples had a high abundance of *Corynebacterium* and *Staphylococcus* (profile IV). In contrast to 16S rRNA analysis, only eight obligate anaerobes were cultured from HS samples and *Porphyromonas* in specific could not be cultured at all (table 2). Although obligate anaerobes were identified in previous culture-based studies^37–41,55^, this was rather limited in our study, indicating a potential underestimation of anaerobic bacteria, associated with sub-optimal culturing conditions for anaerobes^56^.

Distance-based redundancy analysis did not reveal an association of the Hurley classifications with the bacterial composition, as confirmed by the differential abundance analysis. Moreover, no specific bacterial HS profile (Figure1E) could be linked to each Hurley stage. This suggests that the bacterial composition is not associated with the disease severity defined by the Hurley classification system. However, *Enterococcus* and *Streptococcus* were cultured more at Hurley stage III. This correlation of *Streptococcus* with Hurley stage III was also reported previously^46^. Although these bacteria have a commensal relationship with the human body, some species are considered opportunistic pathogens^57,58^. For example, *S. anginosus* is part of the *Streptococcus anginosus* group, known for their ability to cause abscesses^59^. In addition, *P. aeruginosa* and *M. morganii* were isolated only in Hurley III samples. It is therefore possible that at a higher disease severity, the lesions of patients become colonized with these opportunistic pathogens. This is supported by the negative correlation of disease severity with the abundance of skin commensals by Naik *et al.*^47^. Furthermore, the HS profile clustering farthest away (profile I) from the healthy samples (profile IV) did not contain any of the Hurley I samples, whereas all samples from noduli were present in the profile closest to the healthy cluster. This suggests that the lesion type does influence the bacterial profile, but that the lesion activity at the moment of sampling, the presence of puss and depth of the lesions needs to be taken into account.

Both HS patients with and without additional IBD were included in this study, but the variable IBD had no significant effect on the microbiome composition of HS patients. This was confirmed by hierarchical clustering, in which IBD was not significantly associated with the subgrouping of the samples. This is the first time being shown that a comorbidity with IBD does not have a significant effect on the bacterial composition of HS lesions.

## Conclusion

Lesions of Hidradenitis Suppurativa patients have a distinct bacterial composition compared to the skin of healthy individuals. In here, a bacterial imbalance is observed in which mainly *Escherichia-Shigella* and *Porphyromonas* take over the presence of *Staphylococcus* and *Corynebacterium*. Although no clear microbial profile could be linked to each of the Hurley stages, it seems that the bacterial community in skin lesions is dominated by opportunistic pathogens when patients have a higher disease severity. Furthermore, we here show that a comorbidity with IBD does not influence the bacterial composition of HS lesions.

---

**Figure S1:**
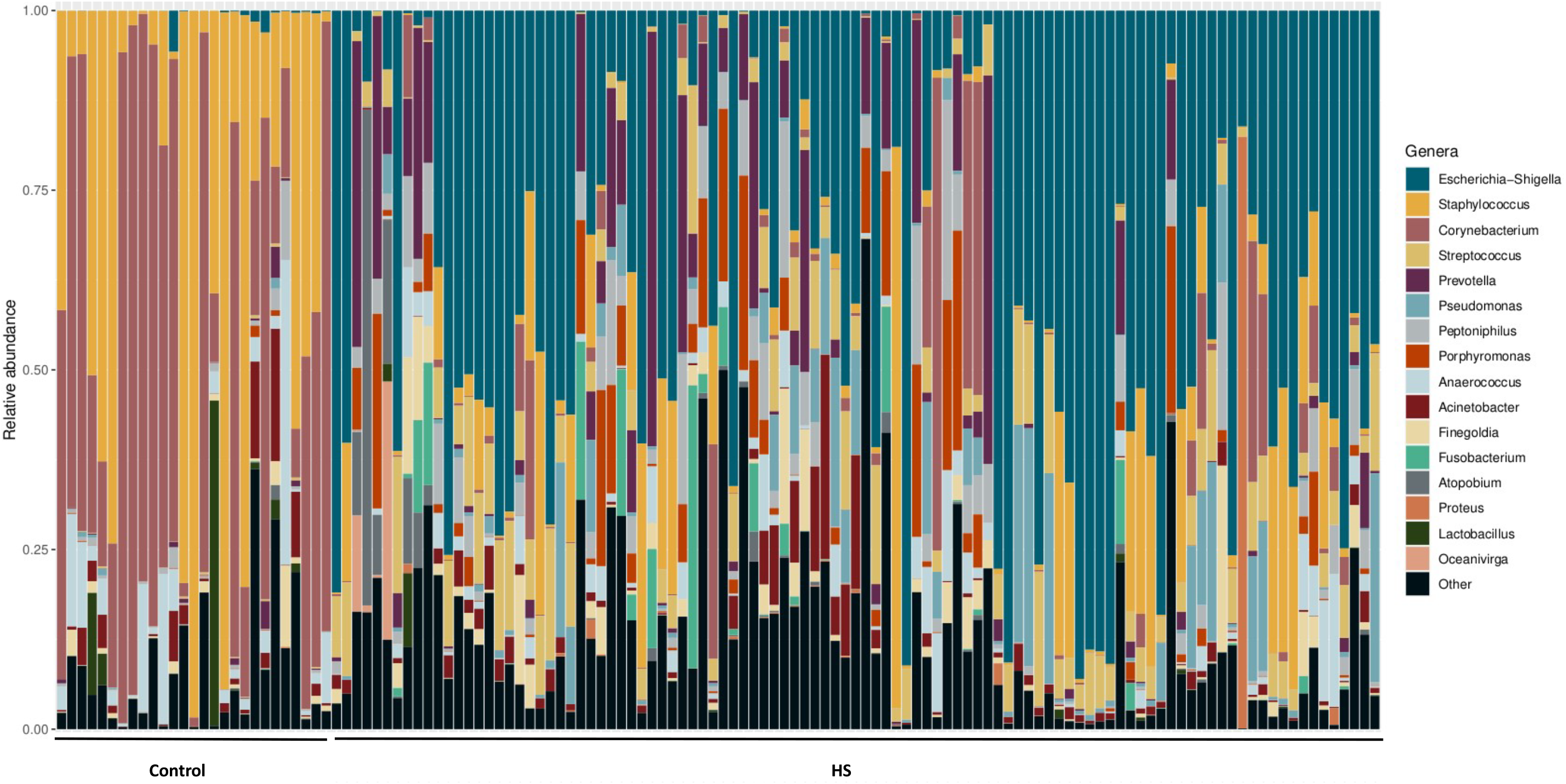
Taxonomic barplot. The relative abundance of the genera in each sample is shown. Only genera with a minimal relative abundance of 15% in at least one sample are shown, the other genera are grouped together in “Other”.

## References

1. Van der Zee, H. H., Laman, J. D., Boer, J. & Prens, E. P. Hidradenitis suppurativa: Viewpoint on clinical phenotyping, pathogenesis and novel treatments. Experimental Dermatology vol. 21 735–739 Preprint at 10.1111/j.1600-0625.2012.01552.x (2012).

2. Sabat, R. et al. Hidradenitis Suppurativa. Nature reviews: disease primers 6, (2020).

3. Morss-Walton, P. C., Kimball, A. B. & Porter, M. L. 2 - Hidradenitis Suppurativa Epidemiology. in (eds. Shi, V. Y., Hsiao, J. L., Lowes, M. A. & Hamzavi, I. H. B. T.-A. C. G. to H. S.) 10–17 (Elsevier, 2022). 10.1016/B978-0-323-77724-7.00002-4.

4. Principi, M. et al. Hydradenitis suppurativa and inflammatory bowel disease: An unusual, but existing association. World Journal of Gastroenterology 22, 4802–4811 (2016).

5. Chen, W.-T. & Chi, C.-C. Association of Hidradenitis Suppurativa With Inflammatory Bowel Disease. JAMA Dermatology 155, 1022–1027 (2019).

6. Zouboulis, C. C. et al. What causes hidradenitis suppurativa ?-15 years after. Experimental dermatology 29, 1154–1170 (2020).

7. Ring, H. C. et al. The follicular skin microbiome in patients with hidradenitis suppurativa and healthy controls. JAMA Dermatology 153, 897–905 (2017).

8. Naik, H. B. et al. Are Bacteria Infectious Pathogens in Hidradenitis Suppurativa? Debate at the Symposium for Hidradenitis Suppurativa Advances Meeting, November 2017. Journal of Investigative Dermatology vol. 139 13–16 Preprint at 10.1016/j.jid.2018.09.036 (2019).

9. Hendricks, A. J., Hsiao, J. L. & Shi, V. Y. 14 - Overview and Comparison of Hidradenitis Suppurativa Management Guidelines. in (eds. Shi, V. Y., Hsiao, J. L., Lowes, M. A. & Hamzavi, I. H. B. T.-A. C. G. to H. S.) 130–144 (Elsevier, 2022). 10.1016/B978-0-323-77724-7.00014-0.

10. Robert, E. et al. Non-surgical treatments for hidradenitis suppurativa: A systematic review. Annales de Chirurgie Plastique Esthetique 62, 274–294 (2017).

11. Revuz, J. Place de la chirurgie dans le traitement de l’hidradénite suppurée. Annales de Dermatologie et de Venereologie 135, 349–350 (2008).

12. Bens, L. et al. Phage Therapy for Hidradenitis Suppurativa: a unique challenge and opportunity for personalized treatment of a complex, inflammatory disease. Clinical and experimental dermatology (2023) doi:10.1093/ced/llad207.

13. Jansen, D. et al. Hidradenitis Suppurativa Patients Exhibit a Distinctive and Highly Individualized Skin Virome. bioRxiv 2023.10.30.564771 (2023) doi:10.1101/2023.10.30.564771.

14. Canoui-Poitrine, F. et al. Clinical characteristics of a series of 302 French patients with hidradenitis suppurativa, with an analysis of factors associated with disease severity. Journal of the American Academy of Dermatology vol. 61 51–57 Preprint at 10.1016/j.jaad.2009.02.013 (2009).

15. Van der Zee, H. H. & Jemec, G. B. New insights into the diagnosis of hidradenitissuppurativa: Clinical presentations and phenotypes. Journal of American Academy of Dermatology 73, S23–S26 (2015).

16. Sayers, E. W. et al. Database resources of the national center for biotechnology information. Nucleic acids research 50, D20–D26 (2022).

17. Pichler, M. et al. A 16S rRNA gene sequencing and analysis protocol for the Illumina MiniSeq platform. MicrobiologyOpen 7, 1–9 (2018).

18. Bolyen E, Rideout JR, Dillon MR, Bokulich NA, Abnet CC, Al-Ghalith GA, Alexander H, Alm EJ, Arumugam M, Asnicar F, Bai Y, Bisanz JE, Bittinger K, Brejnrod A, Brislawn CJ, Brown CT, Callahan BJ, Caraballo-Rodríguez AM, Chase J, Cope EK, Da Silva R, Diener, C. J. Reproducible, interactive, scalable and extensible microbiome data science using QIIME 2. Nature Biotechnology 37, 852–857 (2019).

19. Mcdonald, D. et al. The Biological Observation Matrix (BIOM) format or: how I learned to stop worrying and love the ome-ome. 1–6 (2012).

20. Callahan, B. J. et al. DADA2: High-resolution sample inference from Illumina amplicon data. 13, (2016).

21. Pedregosa, F., Weiss, R. & Brucher, M. Scikit-learn: Machine Learning in Python. 12, 2825–2830 (2011).

22. Mckinney, W. Data Structures for Statistical Computing in Python. 1, 56–61 (2010).

23. Bokulich, N. A. et al. Optimizing taxonomic classification of marker-gene amplicon sequences with QIIME 2’s q2-feature-classifier plugin. 1–17 (2018).

24. Rognes, T., Flouri, T., Nichols, B., Quince, C. & Mahé, F. VSEARCH: a versatile open source tool for metagenomics. 1–22 (2016) doi:10.7717/peerj.2584.

25. Robeson, M. S., et al. RESCRIPt: Reproducible sequence taxonomy reference database management. (2021). doi:10.1371/journal.pcbi.1009581.

26. Pruesse, E. et al. SILVA: a comprehensive online resource for quality checked and aligned ribosomal RNA sequence data compatible with ARB. 35, 7188–7196 (2007).

27. Quast, C. et al. The SILVA ribosomal RNA gene database project: improved data processing and web-based tools. 41, 590–596 (2013).

28. Heidrich, V., Karlovsky, P. & Beule, L. ‘SRS’ R Package and ‘q2-srs’ QIIME 2 Plugin: Normalization of Microbiome Data Using Scaling with Ranked Subsampling (SRS). Applied Sciences 11, (2021).

29. Team, R. C. R: A Language and Environment for Statistical Computing. Preprint at (2022).

30. Bisanz, J. E. qiime2R: Importing QIIME2 artifacts and associated data into R sessions. (2018).

31. McMurdie, P. J. & Holmes, S. phyloseq: An R Package for Reproducible Interactive Analysis and Graphics of Microbiome Census Data. PLOS ONE 8, e61217 (2013).

32. Wickham, H. ggplot2: Elegant Graphics for Data Analysis. Preprint at (2016).

33. Gloor, G. B., Macklaim, J. M. & Fernandes, A. D. Displaying Variation in Large Datasets: Plotting a Visual Summary of Effect Sizes. Journal of Computational and Graphical Statistics 25, 971–979 (2016).

34. Oksanon, J. et al. vegan: Community Ecology Package. (2022).

35. Kolde, R. pheatmap: Pretty Heatmaps. Preprint at (2019).

36. Revuz, J. Hidradenitis suppurativa. Journal of European Academy of Dermatology and Venereology 23, 985–998 (2009).

37. Brook, I. & Frazier, E. H. Aerobic and anaerobic microbiology of axillary hidradenitis suppurativa. Journal of Medical Microbiology 48, 103–105 (1999).

38. Lapins, J., Jarstrand, C. & Emtestam, L. Coagulase-negative staphylococci are the most common bacteria found in cultures from the deep portions of hidradenitis suppurativa lesions, as obtained by carbon dioxide laser surgery. British Journal of Dermatology 140, 90–95 (1999).

39. Sartorius, K., Killasli, H., Oprica, C., Sullivan, A. & Lapins, J. Bacteriology of hidradenitis suppurativa exacerbations and deep tissue cultures obtained during carbon dioxide laser treatment. British Journal of Dermatology 166, 879–883 (2012).

40. Guet-revillet, H. et al. Bacterial Pathogens Associated with Hidradenitis Suppurativa, France. Emerging Infectious Diseases 20, 1990–1998 (2014).

41. Nikolakis, G. et al. Bacterial colonization in hidradenitis suppurativa/acne inversa: A cross-sectional study of 50 patients and review of the literature. Acta Dermato-Venereologica 97, 493–498 (2017).

42. Christensen, G. J. M. & Brüggemann, H. Bacterial skin commensals and their role as host guardians. Beneficial Microbes 5, 201–215 (2014).

43. Grice, E. A. et al. Topographical and temporal diversity of the human skin microbiome. Science 324, 1190–1192 (2009).

44. Dekaboruah, E., Suryavanshi, M. V., Chettri, D. & Verma, A. K. Human microbiome: an academic update on human body site specific surveillance and its possible role. Archives of Microbiology 202, 2147–2167 (2020).

45. Schneider, A. et al. Loss of microbial diversity and body site heterogeneity in individuals with Hidradenitis Suppurativa. Journal of investigative dermatology 140, (2019).

46. Guet-Revillet, H. et al. The Microbiological Landscape of Anaerobic Infections in Hidradenitis Suppurativa: A Prospective Metagenomic Study. Clinical Infectious Diseases 65, 282–291 (2017).

47. Naik, H. B., Jo, J. H., Paul, M. & Kong, H. H. Skin Microbiota Perturbations Are Distinct and Disease Severity–Dependent in Hidradenitis Suppurativa. Journal of Investigative Dermatology 140, 922–925 (2020).

48. Skowron, K. et al. Human Skin Microbiome: Impact of Intrinsic and Extrinsic Factors on Skin Microbiota. Microorganisms 9, (2021).

49. Rhoads, D. D., Cox, S. B., Rees, E. J., Sun, Y. & Wolcott, R. D. Clinical identification of bacteria in human chronic wound infections: culturing vs. 16S ribosomal DNA sequencing. BMC Infectious Diseases 12, 321 (2012).

50. Williams, S. C., Frew, J. W. & Krueger, J. G. A systematic review and critical appraisal of metagenomic and culture studies in hidradenitis suppurativa. Experimental Dermatology 1–10 (2020) doi:10.1111/exd.14141.

51. Janda, J. M. & Abbott, S. L. 16S rRNA Gene Sequencing for Bacterial Identification in the Diagnostic Laboratory: Pluses, Perils, and Pitfalls. Journal of Clinical Microbiology 45, 2761–2764 (2007).

52. Moran, G. et al. Methicillin-Resistant S. aureus Infections among Patients in the Emergency Department. The new england journal of medicine 335, 666–674 (2006).

53. Ring, H. C. et al. The microbiome of tunnels in hidradenitis suppurativa patients. Journal of the European Academy of Dermatology and Venereology 33, 1775–1780 (2019).

54. Hispán, P. et al. Identification of bacterial DNA in the peripheral blood of patients with active hidradenitis suppurativa. Archives of Dermatological Research 312, 159–163 (2020).

55. Haskin, A., Fischer, A. H. & Okoye, G. A. Prevalence of Firmicutes in Lesions of Hidradenitis Suppurativa in Obese Patients. JAMA Dermatology 152, 1276–1278 (2016).

56. Hanišáková, N., Vítězová, M. & Rittmann, S. K.-M. R. The Historical Development of Cultivation Techniques for Methanogens and Other Strict Anaerobes and Their Application in Modern Microbiology. Microorganisms 10, (2022).

57. Krawczyk, B., Wityk, P., Gałęcka, M. & Michalik, M. The Many Faces of Enterococcus spp.-Commensal, Probiotic and Opportunistic Pathogen. Microorganisms 9, (2021).

58. Krzyściak, W., Pluskwa, K. K., Jurczak, A. & Kościelniak, D. The pathogenicity of the Streptococcus genus. European Journal of Clinical Microbiology & Infectious Diseases 32, 1361–1376 (2013).

59. Belko, J., Goldmann, D. A., Macone, A. & Zaidi, A. K. M. Clinically significant infections with organisms of the Streptococcus milleri group. Pediatric Infectious Disease Journal 21, 715–723 (2002).

